# The 55K SNP-Based Exploration of QTL for Spikelet Number Per Spike in a Tetraploid Wheat (*Triticum turgidum* L.) Population: Chinese Landrace ‘Ailanmai’ × Wild Emmer

**DOI:** 10.1101/2020.10.21.348227

**Authors:** Ziqiang Mo, Jing Zhu, Jiatai Wei, Jieguang Zhou, Qiang Xu, Huaping Tang, Yang Mu, Mei Deng, Qiantao Jiang, Yaxi Liu, Guoyue Chen, Jirui Wang, Pengfei Qi, Wei Li, Yuming Wei, Youliang Zheng, Xiujin Lan, Jian Ma

## Abstract

The spikelet number per spike (SNS) is a primary factor determining wheat yield. Common wheat breeding reduces the genetic diversity among elite germplasm resources, leading to a detrimental effect on future wheat production. It is, therefore, necessary to explore new genetic resources for SNS to increase wheat yield. A tetraploid landrace ‘Ailanmai’ × wild emmer wheat recombinant inbred line (RIL) population was used to construct a genetic map using the wheat 55K single nucleotide polymorphism (SNP) array. The linkage map containing 1150 bin markers with a total genetic distance of 2411.8 cM was obtained. Based on phenotypic data from eight environments and best linear unbiased prediction (BLUP) values, five quantitative trait loci (QTL) for SNS were identified, explaining 6.71-29.40% of the phenotypic variation. Two of them, *QSns.sau-AM-2B.3* and *QSns.sau-AM-3B.2*, were detected as the major and novel QTL. Their effects were further validated in two additional F_2_ populations using tightly linked Kompetitive Allele-Specific PCR (KASP) markers. Potential candidate genes within the physical intervals of corresponding QTL were predicted to participate in inflorescence development and spikelet formation. Genetic associations between SNS and other agronomic traits were also detected and analyzed. This study demonstrates the feasibility of wheat 55K SNP array developed for common wheat in genetic mapping of tetraploid population and shows the potential application of wheat related species in wheat improvement programs.

## Introduction

Wheat (*Triticum* sp.) is among the most widely planted crops globally that provides about 20% of the calories consumed by humans (http://www.fao.org/faostat). Breeding high-yield wheat cultivars is a sustainable approach to meet the demands of a growing human population. Wheat yield is determined by spike number per unit area (SNPA), kernel number per spike (KNS), and kernel weight (KW). KNS is jointly determined by the spikelet number per spike (SNS) and the kernel number per spikelet (KNL). SNS depends on the number of lateral spikelets produced before the spike meristem transitions to the terminal spikelet. It is a complex quantitative trait greatly affected by genetic and environmental factors (Gao et al., 2019; Kuzay et al., 2019). Therefore, dissecting the genetic mechanism of SNS at the QTL level can provide more insights into the role of SNS in yield formation (Ma et al., 2019a; Yao et al., 2019; Chen et al., 2020; Li et al., 2021).

A few genes related to spikelet number and morphology have been reported. For example, the *WHEAT FRIZZY PANICLE* (*WFZP*) gene encodes a member of the APETALA2/ethylene response transcription factor (AP2/ERF) family, which influences the supernumerary spikelet trait in common wheat (Dobrovolskaya et al., 2015). Regulatory expression of the two pivotal flowering genes, *Photoperiod-1* (*Ppd-1*) and *FLOWERING LOCUS T* (*FT*), increases the number of fertile spikelets (Boden et al., 2015). The inflorescence growth rate and the development of paired spikelets are controlled by the *TEOSINTE BRANCHED1* (*TB1*) expression level (Dixon et al., 2018). Similarly, the ortholog of the rice *APO1, WHEAT ORTHOLOG OF APO1* (*WAPO1*) affects the spikelet number (Kuzay et al., 2019). Overexpression of *TaTFL-2D* increases the number of spikelets and florets (Wang et al., 2017).

More loci for SNS should thus be identified and utilized owing to its important role in yield formation. Nonetheless, common wheat breeding reduces the genetic diversity among elite germplasm resources, thus adversely affecting wheat yield (Cavanagh et al., 2013; Sthapit et al., 2020). Notably, the numerous genetic resources for agronomic traits and disease resistance from wheat related species may effectively solve future wheat production challenges (Zaïm et al., 2017; El Haddad et al., 2021). As the progenitor of modern cultivated tetraploid and hexaploid wheat, wild emmer wheat (*T*. *turgidum* subsp. *dicoccoides*) is the secondary gene pool of common wheat. It is resistant to various diseases and possesses important agronomic traits, thereby playing a potentially significant role in wheat breeding (Xie and Nevo, 2008; Yu et al., 2020). For example, *Yr15* derived from wild emmer wheat confers resistance to yellow rust (Klymiuk et al., 2018). The reduced-function allele *GNI-A1* increases the number of fertile florets per spikelet (Sakuma et al., 2019). Cognizant of this, exploiting the genetic resources of wild emmer wheat is crucial in solving the existing bottleneck in wheat breeding. Wheat landraces are also primary materials for wheat improvement. They contain unique gene resources preserved under long-term natural selection and human intervention with strong adaptability to local environmental conditions and high production potential (Lopes et al., 2015; Crossa et al., 2016).

In the current research, a high-quality genetic map was constructed based on a RIL population developed from the cross between a Chinese landrace ‘Ailanmai’ and wild emmer using the wheat 55K SNP array to identify QTL for SNS, and major and novel QTL were validated in the different genetic backgrounds by KASP markers.

## Materials and Methods

### Plant Materials

A tetraploid wheat population (AM population) containing 121 F_8_ recombinant inbred lines (RILs, including two parental lines) derived from a Chinese landrace ‘Ailanmai’ (AL, *T*. *turgidum* L. cv. Ailanmai, 2n=28, AABB) and a wild emmer accession (LM001, *T*. *turgidum* subsp. *dicoccoides*) were used in this study. AL is a durum wheat landrace native to Sichuan Province, China. It has the advantages of dwarf, multiple floret and strong environmental adaptability (Liu et al., 1999). AM RIL population was developed aiming at identifying favorable alleles from AL and LM001 to speed up their breeding utilization. The spike morphology of the parents and representative lines are shown in Figure 1. Major and novel SNS QTL identified in the AM population were validated in two populations including LM001 × PI193877 (*T. turgidum* ssp. *dicoccon*) (139 F_2_ lines) and AL × AS2268 (*T. carthlicum* Nevski) (100 F_2_ lines). The above plant materials were preserved by Triticeae Research Institute, Sichuan Agricultural University.

**Figure 1.**
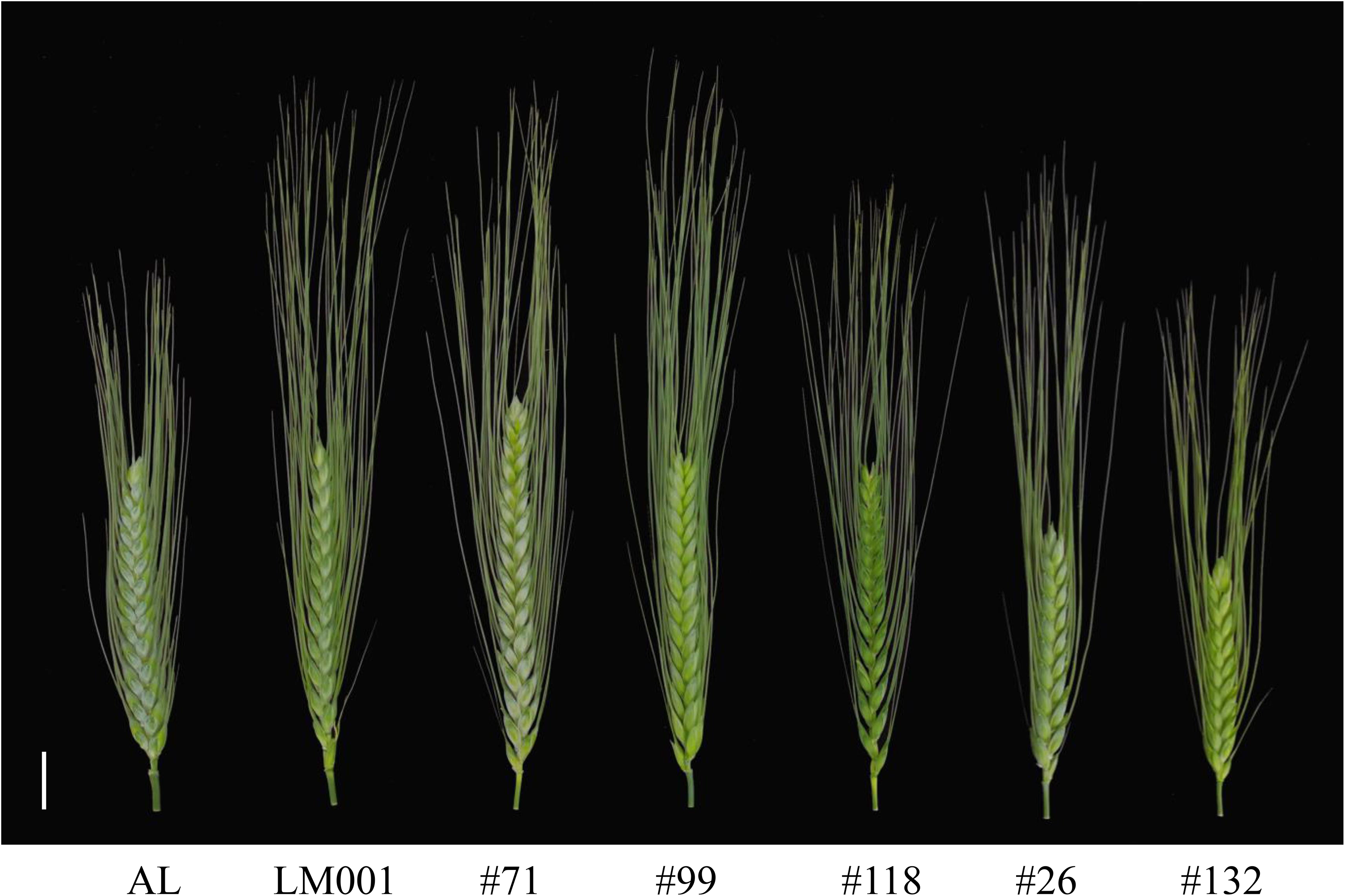
The spike morphology of the parents and representative lines from the AM population. (scale bar = 2 cm).

### Trait Measurement and Data Analysis

Trait measurements of AM population were performed in eight environments: Chongzhou (103° 38’ E, 30° 32’ N) in 2017, 2018, 2019, 2020, and 2021 (E_1_, E_2_, E_3_, E_4_, and E_7_), Wenjiang (103° 51’ E, 30° 43’ N) in 2020 and 2021 (E_5_ and E_8_), and Ya’an (103° 0’ E, 29° 58’ N) in 2020 (E_6_). Field experiments adopted a randomized complete block design, and were managed following the conventional practices of wheat production. The plot for a single line was space-planted (0.1-m) in a single 1.5-m row with a 0.3-m of row spacing. The agronomic traits were measured using the methods described by Liu et al. (2020a). Spikelet number per spike (SNS) was measured by counting the spikelet number of the main-spike per plant; plant height (PH) was determined by the distance from the soil surface to the tip of the main-spike (excluding awns) per plant; spike length (SL) was measured by the length from the base to the tip of the main-spike (excluding awns) per plant; kernel number per spike (KNS) was counted as the kernel number of the main-spike per plant; spike density (SD) was calculated by dividing the SNS by SL; thousand kernel weight (TKW) was calculated as 10 times the average weight of 100 seeds with three repetitions in each line. Anthesis date (AD) was the number of days between sowing and half of the plants flowering in each line (Ma et al., 2019a). Spike extension length (SEL) was measured as the distance between the base of the main-spike and the petiole of the flag leaf per plant (Li et al., 2020). At least four disease-free plants with consistent growth status were selected for measurements. The environmental information of surveyed agronomic traits of AM population is listed in Supplementary Table 1. Two F_2_ validation populations were grown in Chongzhou in 2021 and disease-free plants were selected to measure SNS.

The IBM SPSS Statistics 25 (Armonk, NY, USA) was used to analyze the phenotypic data, including Pearson’s correlation, frequency distribution, standard error, and the Student’s *t*-test (*P* < 0.05). The broad-sense heritability (*H*^*2*^) and best linear unbiased prediction (BLUP) of agronomic traits from multiple environments were calculated using SAS version 8.0 (Cary, North Carolina).

### Construction of the Genetic Map

High-quality genomic DNA was isolated from fresh leaves using the Plant Genomic DNA Kit (Tiangen Biotech, Beijing, China). The DNA of 119 RILs and two parental lines was subsequently genotyped by CapitalBio Technology (Beijing, China) using the wheat 55K SNP array, and RILs were used for linkage map construction.

The genetic map was constructed following the methods described by Liu et al. (2018) and Lin et al. (2021). Firstly, the Poly High-Resolution markers from A and B genomes showing a minor allele frequency (≥ 0.3) among the mapping population were retained for subsequent analysis. Secondly, the BIN function of QTL IciMapping 4.1 (Meng et al., 2015) was used to analyze the remaining markers based on their segregation patterns in the mapping population, with the parameters ‘Distortion value’ and ‘Missing rate’ being set as 0.01 and 20%, respectively. A single marker with the lowest ‘Missing rate’ in each bin (bin marker) was selected to construct genetic maps. Finally, the bin markers were grouped and sorted using the Kosambi mapping function in IciMapping 4.1 with the logarithm of odds (LOD) ≥ 6 after preliminary analysis of SNP markers with LOD values ranging from 2 to 10. The genetic maps were drawn using MapChart 2.3.2 (Voorrips, 2002). The syntenic relationships between the genetic and physical maps of the bin markers were further presented using the Strawberry Perl 5.24.0.1 (Rothwell, 2019).

### QTL Analysis

QTL analysis was performed using the inclusive composite interval mapping (ICIM) in the Biparental Populations (BIP) module of QTL IciMapping 4.1. The LOD threshold was set as 3.4 based on a test of 1000 permutations in IciMapping which was used to determine the threshold that corresponded to a genome-wide false discovery rate of 5% (*P* < 0.05), and loci detected in only single environment were removed. The QTL-by-Environment interaction of SNS trait was analyzed using the function of multi-environment trials (MET-ADD) in IciMapping (Step = 1 cM, PIN = 0.001, and LOD ≥ 3.4). Epistatic interaction of SNS QTL was also identified using the multi-environmental trials (MET-EPI) pattern of IciMapping. Based on the initial QTL mapping results, we developed new KASP markers to densify the genetic map and narrow the mapping interval. The genomic DNA of two parental lines was collected and genotyped by CapitalBio Technology (Beijing, China) using the wheat 660K SNP array. Polymorphic SNPs detected in initial mapping region of two parental lines were converted to KASP markers to genotype the mapping population and integrated into the genetic map.

QTL identified in at least five environments (including BLUP) and explained more than 10% of the phenotypic variation were regarded as major ones, while those having common flanking markers were treated as a single one. QTL was then named following the Genetic Nomenclature provision (http://wheat.pw.usda.gov/ggpages/wgc/98/Intro.htm). In the naming system, ‘Sau’ represented ‘Sichuan Agricultural University’. The RILs’ name ‘AM’ was also added to the QTL’s name to distinguish them from others reported previously.

Sequences of the markers were blasted against the durum wheat (Svevo) (Maccaferri et al., 2019), wild emmer (Zavitan; v2.0) (Zhu et al., 2019), and common wheat genotype ‘Chinese Spring’ (CS; v2.1) (Zhu et al., 2021) genomes to identify their physical positions. The Svevo and Zavitan genomes were used when considering the physical positions of QTL detected from AL and LM001, respectively. Predicted genes mapped between the flanking markers and their function annotations and expression patterns were obtained from the Triticeae Multi-omics Center (http://202.194.139.32/).

### QTL Validation

The tightly linked SNP markers of major and novel SNS QTL were subjected to develop KASP markers. KASP primers were designed and applied following the method of Tan et al. (2017). Each validation population was divided into two haplotype groups (with homozygous alleles from any parent) according to their alleles present in parents and progenies. Phenotypic differences between the two groups were analyzed using Student’s *t*-test (*P* < 0.05).

## Results

### SNP Markers and Genetic Map

As suggested by the Affymetrix Company, the high probability probes from the Poly High-Resolution group only were reserved. Thus, 8724 SNPs with minor allele frequency (≥ 0.3) among the mapping population were retained for following analysis. Subsequently, 823 markers from the D genome were rejected. Finally, 7901 SNP markers were classified into 1192 bins, and a single marker with the lowest ‘Missing rate’ in each bin (bin marker) was screened to construct the genetic map. The linkage analysis results revealed that 1150 bin markers were mapped on 15 linkage groups. One linkage group was constructed for each chromosome except chromosome 3B, which had two (Table 1 and Supplementary Table 2). Based on the bin information, 6537 mapped markers (including 1150 bin markers and 5387 co-segregated markers) were integrated into the genetic map with a total length of 2411.8 cM (Table 1 and Supplementary Table 3). The length of genetic map in each linkage group ranged between 49.7 cM (3B1) and 225.1 cM (5A). The average distance of bin markers ranged between 1.09 cM (4B) and 3.64 cM (3B2), with an average density of one bin marker per 2.10 cM. The number of mapped markers ranged between 37 on chromosome 3B1 and 944 on chromosome 2A (Table 1 and Supplementary Table 3). Notably, 56% and 44% of the mapped markers were distributed on A and B subgenomes, respectively.

**Table 1.** Information of SNP markers in the constructed genetic map

| **Chromosome** | **Group** | **Number of**<br>**bin markers** | **Number of**<br>**mapped markers** | **Length**<br>**(cM)** | **Density**<br>**(cM/bin)** |
| --- | --- | --- | --- | --- | --- |
| 1A | 1 | 101 | 586 | 153.3 | 1.52 |
| 1B | 1 | 69 | 488 | 149.3 | 2.16 |
| 2A | 1 | 66 | 944 | 161.8 | 2.45 |
| 2B | 1 | 92 | 424 | 183.7 | 2.00 |
| 3A | 1 | 99 | 517 | 161.0 | 1.63 |
| 3B | 1 | 17 | 37 | 49.7 | 2.92 |
|  | 2 | 27 | 174 | 98.4 | 3.64 |
| 4A | 1 | 80 | 282 | 199.9 | 2.50 |
| 4B | 1 | 77 | 422 | 83.7 | 1.09 |
| 5A | 1 | 92 | 298 | 225.1 | 2.45 |
| 5B | 1 | 121 | 674 | 207.5 | 1.72 |
| 6A | 1 | 60 | 636 | 167.0 | 2.78 |
| 6B | 1 | 58 | 160 | 162.5 | 2.80 |
| 7A | 1 | 92 | 395 | 214.2 | 2.33 |
| 7B | 1 | 99 | 500 | 194.8 | 1.97 |
| A genome | 7 | 590 | 3658 | 1282.2 | 2.17 |
| B genome | 8 | 560 | 2879 | 1129.6 | 2.02 |
| Total | 15 | 1150 | 6537 | 2411.8 | 2.10 |

### Comparison of Genetic and Physical Maps

Considering that the parent accessions AL and LM001 were durum wheat and wild emmer wheat, respectively, the sequences of the 6537 mapped markers were blasted against durum wheat and wild emmer genomes to identify their physical positions (Supplementary Table 3). The genetic chromosomal locations of the 1150 bin markers were compared with their physical chromosomal locations in the durum wheat genome. Among them, 1087 (94.52 %) markers showed good consistencies (Figure 2a), and 63 (5.48%) markers had inconsistent chromosomal locations. In the wild emmer genome, the genetic chromosomal locations of 71 (6.17%) markers were inconsistent with physical chromosomal locations, while the remaining markers (1079, 93.83%) were well matched (Figure 2b). Notably, 1026 (89.22%) bin markers had consistent chromosomal locations in genetic map, durum wheat genome and wild emmer genome. Similarly, the genetic maps of the 6537 mapped markers were compared with their physical maps (Supplementary Table 3). 6242 (95.49%) and 6249 (95.59%) markers showed consistent chromosomal locations in the durum wheat and wild emmer genomes, respectively, while 6031 (92.26%) markers had consistent chromosomal locations in genetic map and two physical maps.

**Figure 2.**
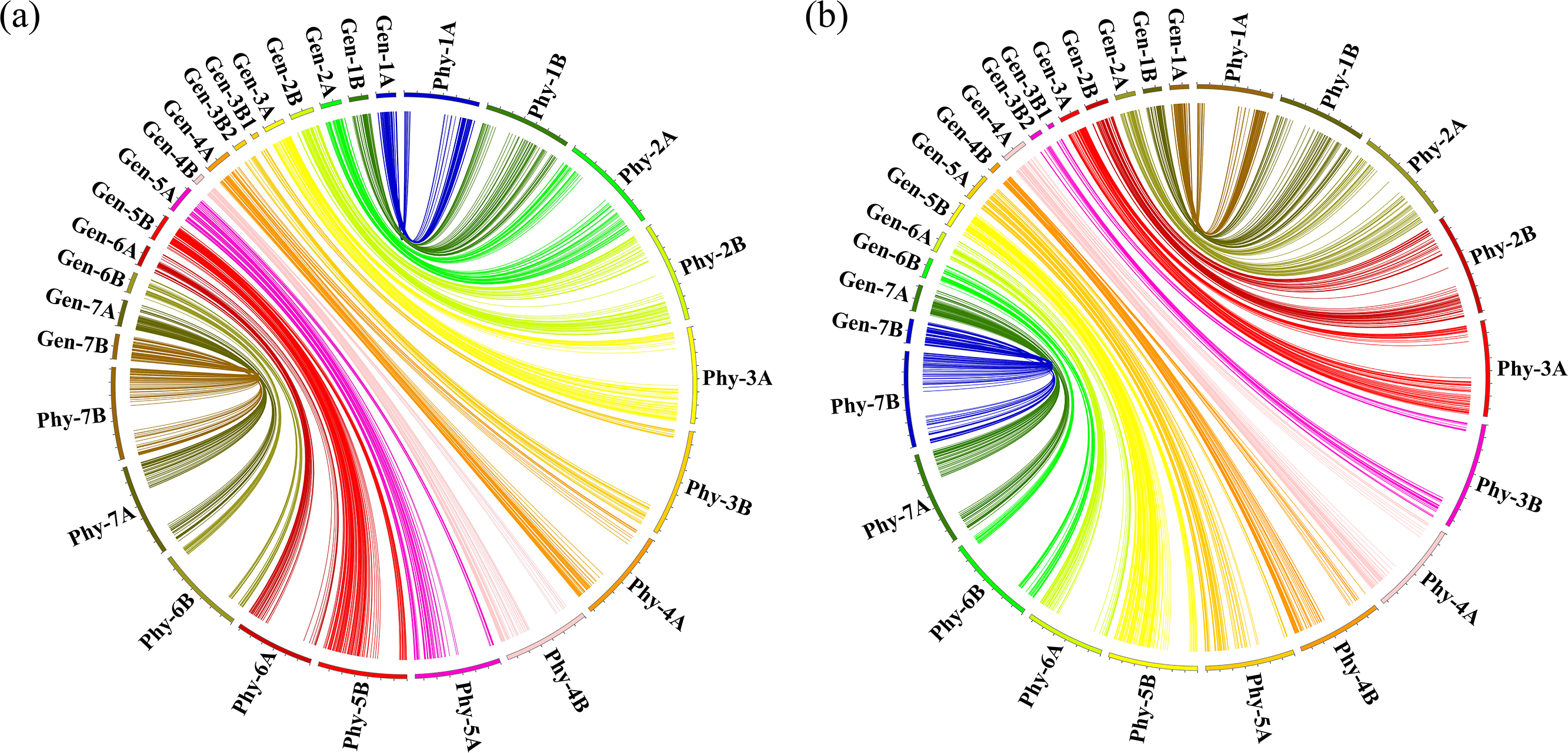
The syntenic relationships between the genetic and physical maps of bin markers. Gen-1A to Gen-7B represented the 15 chromosomal genetic maps used in the current study; Phy-1A to Phy-7B represented the 14 chromosomal physical maps of (**a**) durum wheat and (**b**) wild emmer genomes.

### Phenotypic Variation

The SNS of AL and LM001 in multiple environments ranged between 25.00 and 27.00, and 23.33 and 26.00, respectively (Table 2). That of AM RILs ranged between 15.33 and 34.00. There were no significant differences in SNS between the two parental lines. However, continuous variation and bidirectional transgressive segregation was observed in the RIL population. These results suggested that it was feasible to analyze the SNS loci in the current population.

**Table 2.** Phenotypic data and heritability (*H*^*2*^) of spikelet number per spike (SNS) for the AM population in multiple environments

| Environment | Parents | | AL × LM001 (AM) | | | |
| --- | --- | --- | --- | --- | --- | --- |
| | AL | LM001 | Min-Max | Mean | SD | $H^2$ |
| E<sub>1</sub> | 25.00 | 24.33 | 20.00-31.67 | 25.96 | 2.56 | 0.95 |
| E<sub>2</sub> | 26.00 | 23.33 | 15.33-33.00 | 24.41 | 3.22 | 0.97 |
| E<sub>3</sub> | 26.33 | 26.00 | 17.00-31.25 | 24.81 | 2.80 | 0.96 |
| E<sub>4</sub> | 26.00 | 25.40 | 18.00-32.00 | 26.21 | 2.87 | 0.96 |
| E<sub>5</sub> | 27.00 | 24.60 | 18.50-31.00 | 24.99 | 2.42 | 0.95 |
| E<sub>6</sub> | 25.80 | 25.40 | 20.00-34.00 | 26.76 | 2.69 | 0.96 |
| E<sub>7</sub> | 23.40 | 24.25 | 15.75-31.00 | 23.68 | 2.62 | 0.95 |
| E<sub>8</sub> | 25.44 | 24.67 | 18.50-30.67 | 24.93 | 2.55 | 0.95 |
| BLUP | 25.60 | 24.78 | 18.41-30.10 | 25.22 | 2.12 | 0.93 |

*BLUP* phenotype values based on the best linear unbiased prediction, *SD* standard deviation, $H^2$ the broad-sense heritability

The broad-sense heritability (*H*^*2*^) of SNS ranged between 0.93 and 0.97 in each environment (Table 2), indicating that the SNS trait was mainly controlled by genetic factors. Strong interaction was detected between SNS and environments (*P* < 0.01), with significant correlation coefficients ranging between 0.50 and 0.90 (Figure 3). The frequency distribution of SNS in multiple environments indicated that it was polygenically inherited (Figure 3).

**Figure 3.**
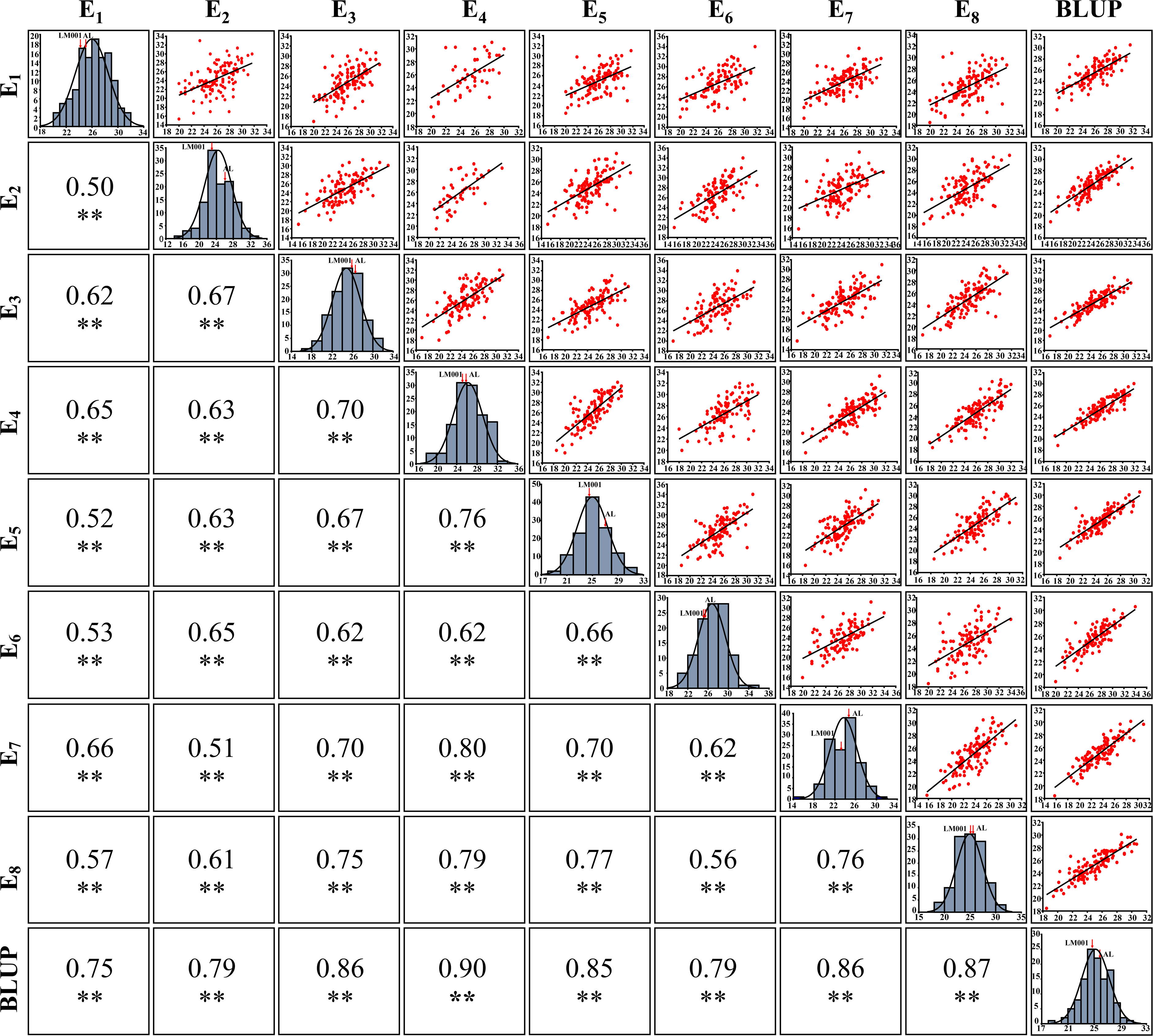
The phenotype, frequency distribution, and correlation coefficients of spikelet number per spike (SNS) in multiple environments. **Significance level at *P* < 0.01.

### Genetic Correlations between SNS and other Agronomic Traits

Pearson’s correlations were analyzed between SNS and seven other agronomic traits based on the BLUP values (Table 3). SNS was significantly and positively correlated with PH, AD, SL, SEL, KNS, and SD. However, SNS was not significantly correlated with TKW.

**Table 3.** Correlations between spikelet number per spike (SNS) and other agronomic traits in the AM population

| Trait | Spikelet number per spike (SNS) |
| --- | --- |
| Plant height (PH) | 0.30** |
| Anthesis date (AD) | 0.34** |
| Spike length (SL) | 0.70** |
| Spike extension length (SEL) | 0.23* |
| Kernel number per spike (KNS) | 0.36** |
| Spike density (SD) | 0.32** |
| Thousand kernel weight (TKW) | 0.14 |

\*Significance level at $P < 0.05$ , \*\*Significance level at $P < 0.01$

### Analysis of QTL Associated with SNS

Five QTL for SNS were identified and distributed on chromosomes 5A (1 QTL), 2B (2), and 3B (2), with LOD scores ranging between 3.55 and 21.49 (Table 4). They explained 6.71-29.40% of the phenotypic variation. *QSns.sau-AM-5A*, *QSns.sau-AM-2B.3*, and *QSns.sau-AM-3B.2* were detected in at least five environments and explained more than 10% of the phenotypic variation, which were treated as major QTL. And the other two QTL were identified as minor QTL.

**Table 4.**
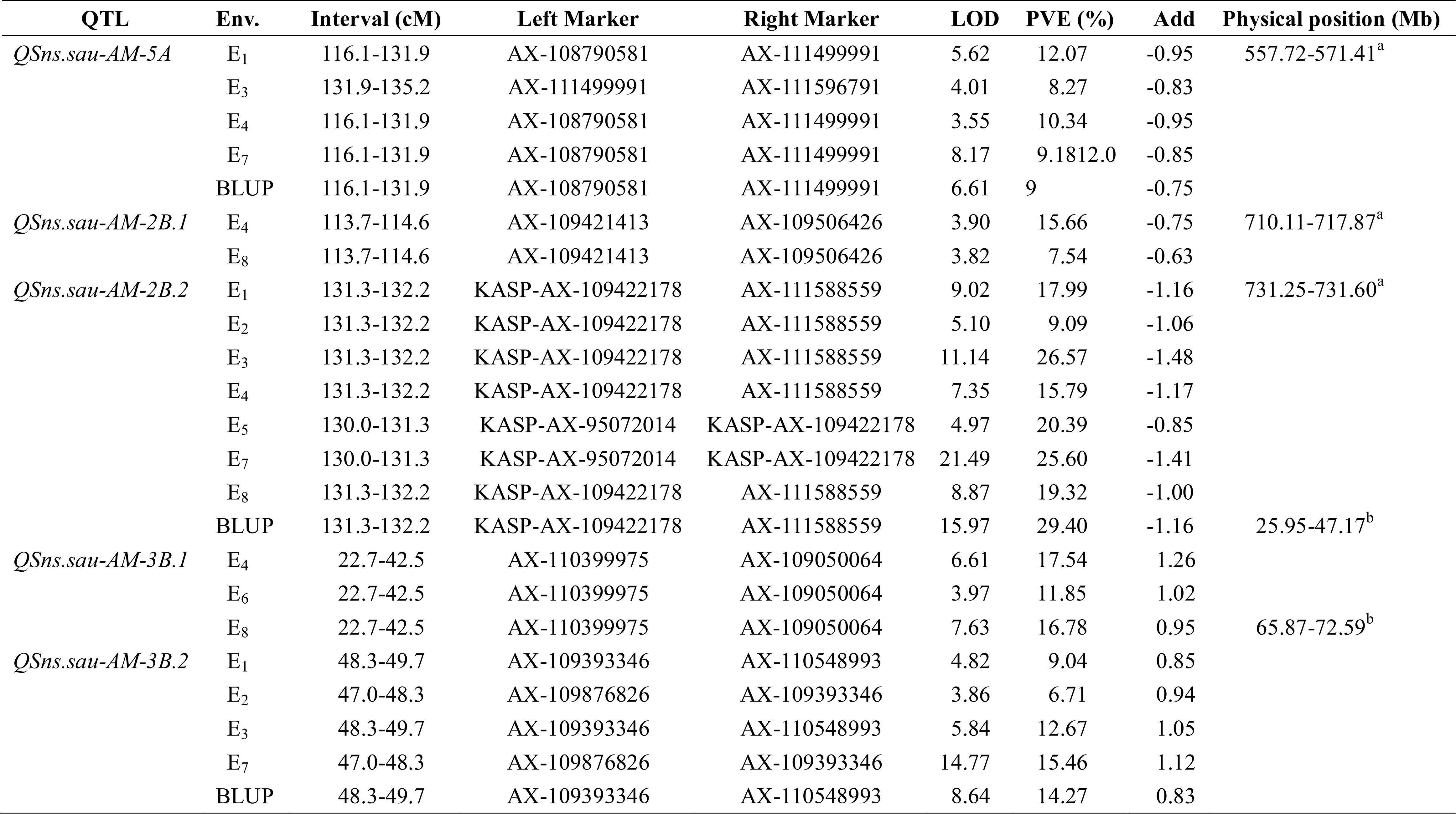
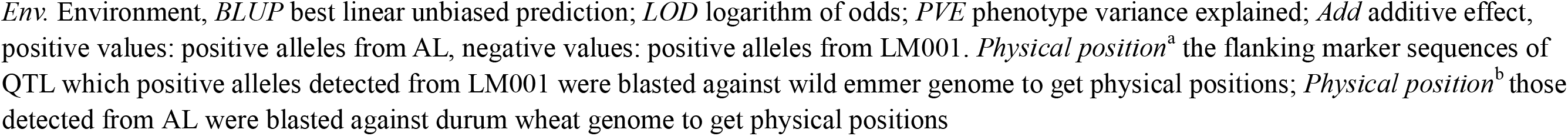
Quantitative trait loci (QTL) for spikelet number per spike (SNS) identified in multiple environments in the AM population

*QSns.sau-AM-5A* was detected in five environments and located in a 19.1 cM region between AX-108790581 and AX-111596791. It accounted for 8.27-12.09% of the phenotypic variation (Table 4). AM population were divided into two haplotype groups based on the genotype of QTL’s flanking markers, and the effect of *QSns.sau-AY-5A.2* increased SNS significantly by 4.54-10.48% (Figure 4a).

**Figure 4.**
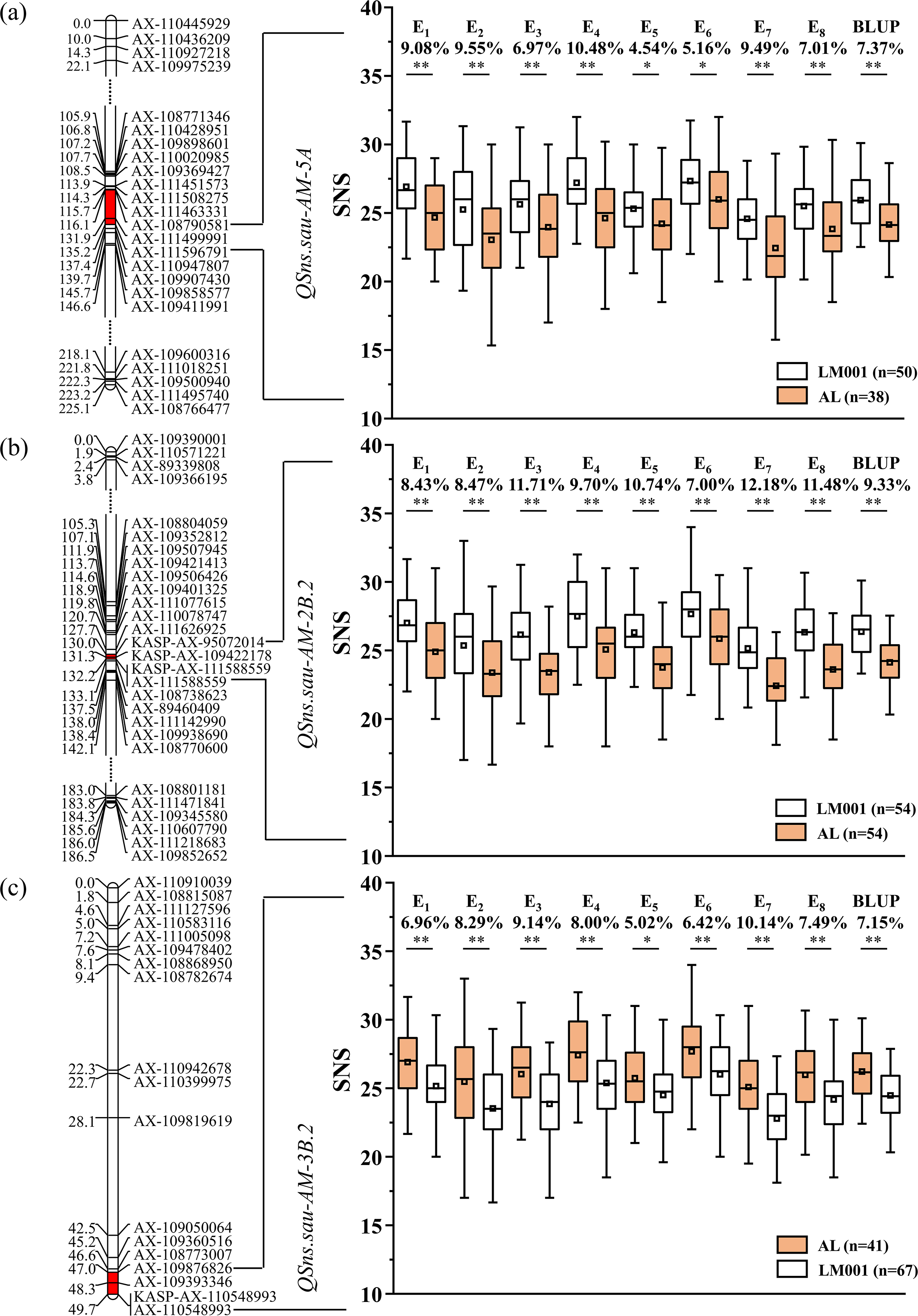
AM RILs were divided into two haplotype groups based on the genotype of flanking markers, and the SNS differences caused by the corresponding QTL were represented. Genetic maps of (**a**) *QSns.sau-AM-5A*, (**b**) *QSns.sau-AM-2B.2*, (**c**) *QSns.sau-AM-3B.2* and their effects. *Significance level at *P* < 0.05, **Significance level at *P* < 0.01.

*QSns.sau-AM-2B.2* was stably identified in all environments, explaining 9.09-29.40% of the phenotypic variation (Table 4). It was firstly mapped between AX-111626925 and AX-111588559 (Figure 4b). AX-95072014 and AX-109422178 were polymorphic SNPs detected in initial mapping interval of two parental lines using the wheat 660K SNP array. They were converted to KASP markers (KASP-AX-95072014 and KASP-AX-109422178) to genotype the mapping population and integrated into the genetic map. Remapping results showed that *QSns.sau-AM-2B.2* was located in a 2.2 cM region between KASP-AX-95072014 and AX-111588559. The additive effect of *QSns.sau-AM-2B.2* was negative, indicating that the increasing SNS allele was contributed by LM001. The effects of *QSns.sau-AM-2B.2* on increasing SNS in single environment ranged from 7.00% to 12.18% (Figure 4b).*QSns.sau-AM-3B.2* was detected in five environments (Table 4). It explained up to 15.46% of the phenotypic variation. The positive allele of *QSns.sau-AM-3B.2* was from AL, and RILs containing the positive allele of *QSns.sau-AM-3B.*2 had a higher SNS than those containing the negative one. The effect of *QSns.sau-AM-3B.2* significantly increased SNS by 5.02-10.14% in single environment (Figure 4c).

The QTL-by-Environment interaction analysis identified 47 QTL (Supplementary Table 4). Five of them were the same as detected in individual environment analysis, indicating that they were stably expressed for SNS. In addition, epistatic interaction were also analyzed, and the results showed that there were no epistasis effects between five SNS QTL detected in the present study (Supplementary Table 5).

### Effects of Major QTL on Increasing SNS

The effects of three QTL (*QSns.sau-AM-5A*, *QSns.sau-AM-2B.2*, and *QSns.sau-AM-3B.2*) on increasing the SNS were further revealed in the mapping population based on the genotypes of the flanking markers (Figure 5). Compared to the haplotype group that carried the negative alleles of three QTL, haplotype groups with the favorable alleles from at least one QTL significantly increased SNS. The haplotype group containing the favorable alleles from three QTL significantly increased SNS compared with those with two or single one, respectively. No significant differences were detected among three haplotype groups with any two positive alleles. *QSns.sau-AM-2B.2* had the largest individual positive effect on SNS. However, there was no significant difference between the individual effect of three QTL on increasing SNS.

**Figure 5.**
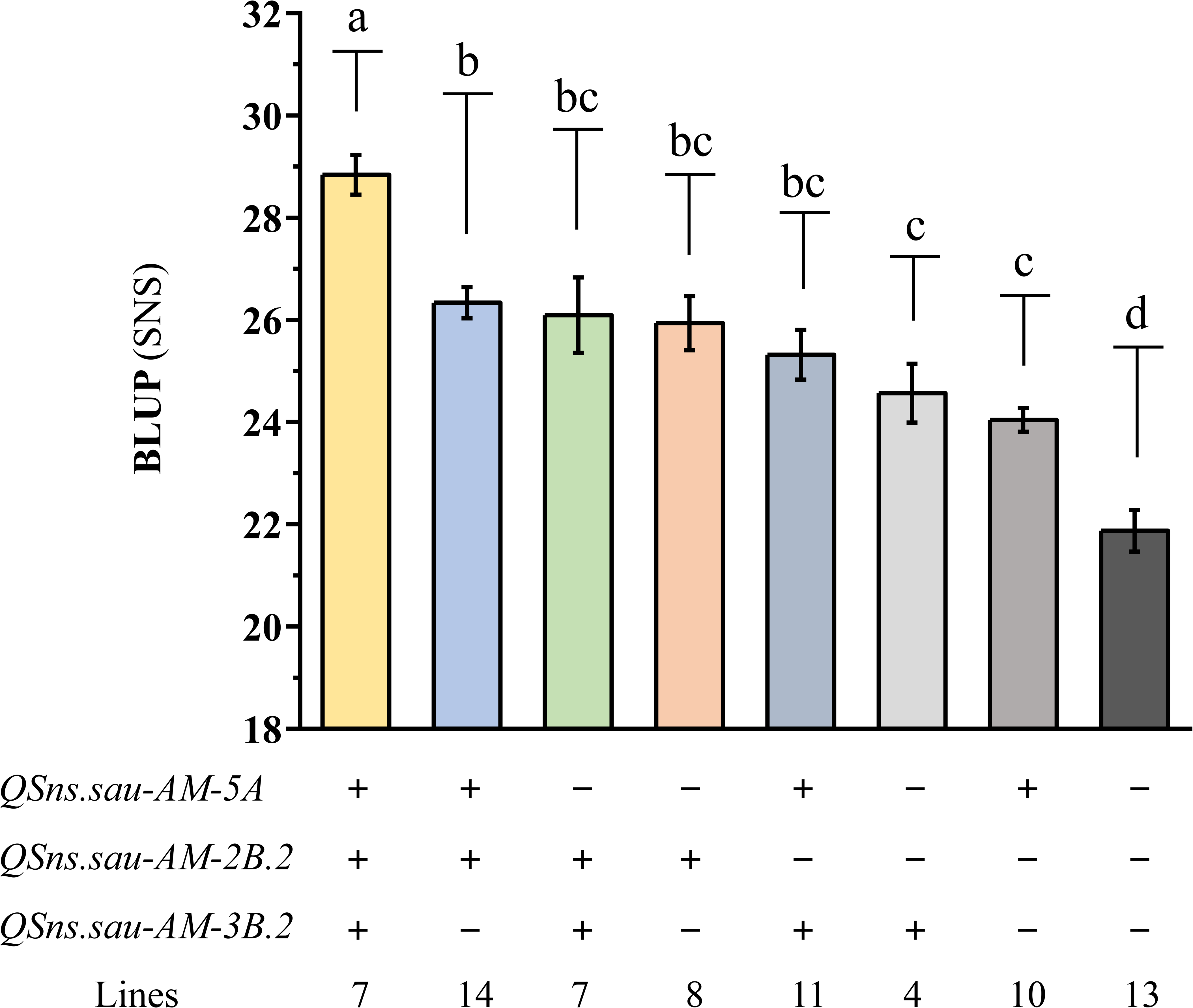
The additive effects of *QSns.sau-AM-5A*, *QSns.sau-AM-2B.2*, and *QSns.sau-AM-3B.2* on increasing SNS. ‘+’ and ‘−’ represented haplotype group with and without the positive allele of the corresponding QTL based on the genotype of flanking markers, respectively. Letters a, b, c, and d represent significant difference among haplotype group.

### Validation of Major and Novel SNS QTL

The SNP markers closely linked to *QSns.sau-AM-2B.2* (AX-111588559) and *QSns.sau-AM-3B.2* (AX-110548993) were converted to KASP markers, to verify the effects of them in two other populations (Supplementary Table 6 and Figure 6). Since the favorable alleles of *QSns.sau-AM-2B.2* and *QSns.sau-AM-3B.2* were from LM001 and AL, respectively, haplotype group with homozygous alleles of LM001 showed a significant increase in SNS by 8.33% (Figure 6a) and those from AL increased SNS significantly by 6.00% (Figure 6b).

**Figure 6.**
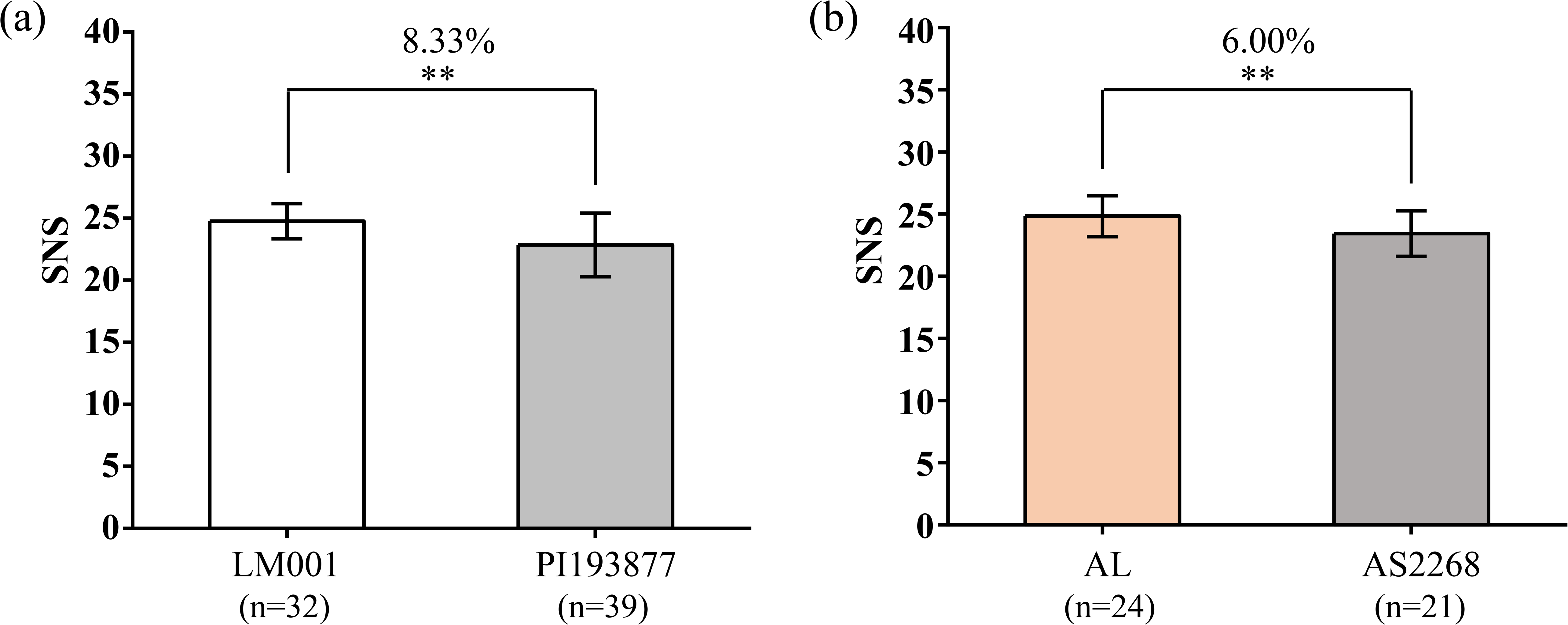
Validation of (**a**) *QSns.sau-AM-2B.2* (KASP-AX-111588559) and (**b**) *QSns.sau-AM-3B.2* (KASP-AX-110548993) in LM001 × PI193877 and AL × AS2268 populations by Student’s *t*-test, respectively. **Significance level at *P* < 0.01.

## Discussion

### Exploration of the SNS Loci from Tetraploid Wheat Species Using the Wheat 55K SNP Array

Previous studies have reported on SNS in tetraploid wheat at the QTL level. Peng et al. (2001) identified four SNS QTL, explaining a total of 49.3% of the phenotypic variation in a wild emmer × durum wheat F_2_ population genotyped using restriction fragment length polymorphisms (RFLP) markers. Genotyping of the durum × cultivated emmer wheat RIL population with the 9 K iSelect SNP chip identified four SNS QTL on chromosomes 1B, 3B, 7A, and 7B (Faris et al., 2014a). In the same time, a linkage map of chromosome 2A generated in the Langdon × Langdon-wild emmer wheat substitution lines population consisting of 23 simple sequence repeat (SSR) markers was used to identify an SNS QTL, *QSpn.fcu-2A*, explaining 19.4% of the phenotypic variation (Faris et al., 2014b). SNS QTL, *QSpn.fcu-1B*, and *QSpn.fcu-5A*, have been identified in a durum × cultivated emmer wheat population genotyped using the 90K iSelect array (Sharma et al., 2019). These reports enhance our understanding of the genetic mechanism of SNS in tetraploid wheat.

The wheat 55K SNP array was developed based on the selection and optimization of the 660K SNP array which was designed based on hexaploid, tetraploid and diploid wheat, and has the advantages of being efficient and genome-specific (Sun et al., 2020). It has been widely used for genotyping and genetic loci identification of agronomic traits and disease resistance in bread wheat (Ren et al., 2018; Ma et al., 2019b; Liu et al., 2020b; Ma et al., 2020; Tu et al., 2021). To the author’s knowledge, this is the first study to adopt the wheat 55K SNP array to construct the genetic linkage map of a tetraploid wheat RIL population. Our results confirm the feasibility of using a 55K SNP array developed for common wheat for genetic mapping of tetraploid wheat, thus proving its broad utilization.

### Comparison of the Major SNS QTL Identified with those of Previous Studies

There was no significant difference for SNS detected between the parental lines in multiple environments (Table 2). The phenotypic data of SNS in the six environments showed approximately normal distribution (Figure 4). Three major and stably expressed QTL for SNS were identified. This phenomenon is attributed to the transgressive inheritance in progenies that result in the bidirectional segregation of SNS. Several genes controlling SNS may inhibit each other in a given parental genotype resulting in the absence of the corresponding phenotype in the parents. However, a single locus produced by genetic recombination between parental genotypes can display a conspicuous phenotype in the progenies without interference from other loci that control the same trait. The major SNS loci in the AM RIL population can thus be identified. This is not uncommon in the past studies of QTL identification based on RIL populations (Liu et al., 2020a; Shang et al., 2020).

Physical locations of known SNS QTL were detected to verify whether major QTL were novel (Supplementary Table 7). In wild emmer genome, *QSns.sau-AM-5A* was physically located on chromosome arm 5AL at 557.72-571.41 Mb, overlapping with *QSps.ccsu-5A.2* (568.96-662.44 Mb) (Kumar et al., 2007) and *QTL1755_5A* (569.50-579.91 Mb) (Peng et al., 2003), indicating that they are likely alleles. *QSns.sau-AM-2B.2* was located at 731.25-731.60 Mb on chromosome arm 2BL in wild emmer genome. Previous studies have detected SNS QTL on chromosome arm 2BS. However, none has been reported on 2BL. For example, *Qspn-2007* was located on chromosome arm 2BS with the closest marker Xgwm429 (Manickavelu et al., 2011). *QFsn.sdau-2B-1* was located on chromosome arm 2BS, explaining up to 28.43% of the phenotypic variation (Deng et al., 2017). Some genes regulating SNS, such as *Ppd-B1* (Mohler et al., 2004) and *WFZP-2B* (Dobrovolskaya et al., 2015), have also been detected on chromosome arm 2BS. In the durum wheat genome, *QSns.sau-AM-3B.2* was located on chromosome arm 3BS at 65.87-72.59 Mb. Similarly, Cui et al. (2012) identified two QTL, *QSpn.WJ.3B.1* and *QSpn.WY.3B.1*, all closely linked to the locus Xgwm566. Azadi et al. (2015) detected three QTL for SNS on chromosome arm 3BL. Comparing the physical positions of three major QTL with those previously reported indicates that *QSns.sau-AY-2B.2* and *QSns.sau-AY-3B.2* are probably novel loci controlling SNS in tetraploid wheat.

### Potential Candidate Genes in the Physical Intervals of *QSns.sau-AM-2B.2*

*QSns.sau-AM-2B.2* was physical located in a 0.35 Mb region of wild emmer genome, and 4 genes and 3 functional orthologs were annotated (Supplementary Table 8a). *QSns.sau-AM-3B.2* was located in a 6.72 Mb interval of durum wheat genome with 62 genes and 38 functional orthologs (Supplementary Table 8b). Since the physical interval of *QSns.sau-AM-2B.2* contains only 3 orthologs, it is valuable to analyze function annotations and expression patterns to predict candidate genes.

*TRIDC2BG075290* encodes a protein kinase which involves in cell signal transduction and plant growth. Previous research reported wheat protein kinase gene *TaSnRK2.9*◻5A was signifcantly correlated with high thousand kernel weight and high kernel per spike (Ur Rehman et al., 2019), suggesting that protein kinase can have a positive impact on agronomic traits. In addition, *Stomatal opening factor 1* (*OST1*, *SnRK2.6*) is a homologous gene of *TRIDC2BG075290* in *Arabidopsis*. *OST1* can promote the expression of *FLOWERING LOCUS C* (*FLC*) to participate in the regulation of flowering by ABA (Wang et al., 2013), which may affect inflorescence development and the formation of spikelets. *TRIDC2BG075310* and *TRIDC2BG075320* all encode SAUR-like auxin-responsive protein. It has been reported that plant SAUR-like auxin-responsive protein family is involved in the growth of roots, hypocotyls, leaves and stamens (Chae et al., 2012; Hou et al., 2013). Expression patterns analyses showed that *TRIDC2BG075290* has higher expression levels in spike and flag leaf (Supplementary Figure 1). *TRIDC2BG075310* has a high expression level throughout the spike development stage, while *TRIDC2BG075320* is only expressed in spike. Therefore, we speculate that these 3 functional orthologs may affect the wheat inflorescence development, which can be further determined by fine mapping and gene cloning in future.

### Phenotypic Correlations of Agronomic Traits

SNS was significantly and positively correlated with PH in the present study (Table 3). This finding was consistent with that of Ajmal et al. (2009) and Wu et al. (2012). We speculate that SL is part of PH, and the taller plants have a long spike with more spikelets (Yao et al., 2011). There was a significant and positive relationship between SNS and AD (Table 3), consistent with the results of Ma et al. (2019a). It is supposed that wheat plants that take longer to flower tend to produce longer spikes with more spikelets (Shaw et al., 2013; Kamran et al., 2014; Boden et al., 2015). These results indicate that AD plays an important role in spikelet formation.

Similarly, there was a strong significant and positive correlation between SNS and SL (Table 3). The increase in SL requires a longer spike development stage combined with optimal temperature and light to promote spikelet germination (Shaw et al., 2013). On the whole, floret fertility determines the KNS in wheat. More spikelets produce more fertile florets, thereby increasing the KNS (Gao et al., 2019; Sakuma et al., 2019). This reason potentially caused the SNS to be significantly and positively correlated with KNS (Table 3). Moreover, SNS was significantly and positively correlated with SD because SD is the ratio of SNS to SL (Table 3).

It is postulated that there is a negative correlation between SNS and TKW (Li et al., 2007; Ma et al., 2019a). The negative correlation is attributed to source limitations during kernel filling or more kernels with lower KW in the spike’s distal positions (Quintero et al., 2018). Previous studies also suggest that SNS is negatively correlated with SEL because of competition in product assimilation between SNS and SEL (Bancal, 2008; Li et al., 2020). However, these conclusions are inconsistent with the findings of this study. Herein, SNS was positively correlated with TKW and SEL (Table 3). It probably indicates that the loci controlling SNS play an individual role without interacting with SEL. Previous studies postulate that SEL has a special function in photosynthesis and nutrients and water storage (Bridgemohan and Bridgemohan, 2014; Ávila-Lovera et al., 2017). Therefore, a longer SEL is more conducive for ventilation and transportation of more nutrients and water to kernels thereby increasing the KW.

These conclusions are vital in comprehensively understanding the complex relationships in agronomic traits, and provide new insights into wheat yield improvement. KNS is one of the primary determinants of grain yield. Therefore, wheat breeding programs should focus on increasing SNS under moderate SD in the present study.

## Conclusion

Herein, the wheat 55K SNP array was successfully used to construct a high-quality genetic map that identified five QTL for SNS in a tetraploid wheat RIL population. The additive effects of the major SNS QTL in increasing SNS were revealed in the mapping population. *QSns.sau-AM-2B.2* and *QSns.sau-AM-3B.2* were major and novel QTL, and their effects were successfully validated in other genetic backgrounds using closely linked KASP markers. The genetic correlations between SNS and other agronomic traits were also evaluated. The novel genetic loci of SNS and tightly linked KASP markers will be valuable in further fine mapping and gene cloning.

## Data Availability Statement

All data generated or analyzed during this study are included in this published article and its supplementary information files, further inquiries can be directed to the corresponding author.

## Author Contributions

ZQM and JZ conducted the study and drafted this manuscript. JTW and JGZ participated in phenotype measurement and data analysis. QX and HPT helped with field work and data analysis. YM, MD, QTJ, and YXL participated in data collection and analysis. GYC, JRW, PFQ, and WL did QTL analysis and manuscript revision. YMW and YLZ discussed the results and revised the manuscript. XJL guided the study and revised the manuscript. JM designed the experiments, guided the entire study, participated in data analysis, wrote and extensively revised this manuscript. All authors participated in the research and approved the final manuscript.

## Supplementary Materials

**Supplementary Figure 1** Expression patterns at the different development stage of 3 functional orthologs in the physical interval of *QSns.sau-AM-2B.2*. (**a**) *TRIDC2BG075290*; (**b**) *TRIDC2BG075310*; (**c**) *TRIDC2BG075320*.

**Supplementary Table 1** Environmental information of surveyed agronomic traits in AM population.

**Supplementary Table 2** The 1150 bin markers were used for QTL mapping.

**Supplementary Table 3** Comparison of the genetic and physical positions of 6537 mapped markers (including bin markers and co-segregated markers).

**Supplementary Table 4** Quantitative trait loci (QTL) detected in the QTL-by-Environment interaction module.

**Supplementary Table 5** Epistasis interaction analysis of QTL for spikelet number per spike (SNS).

**Supplementary Table 6** Details of KASP markers.

**Supplementary Table 7** Comparison of the major quantitative trait loci (QTL) with loci previously reported for spikelet number per spike (SNS).

**Supplementary Table 8** Predicted genes in the physical interval of *QSns.sau-AM-2B.2* and *QSns.sau-AM-3B.2*.

## References

Ajmal, S., Zakir, N., and Mujahid, M. (2009). Estimation of genetic parameters and character association in wheat. J. Agric. Biol. Sci. 1, 15–18. doi: 10.5713/ajas.2011.10456.

Ávila-Lovera, E., Zerpa, A.J., and Santiago, L.S. (2017). Stem photosynthesis and hydraulics are coordinated in desert plant species. New Phytol. 216, 1119–1129. doi: 10.1111/nph.14737.

Azadi, A., Mardi, M., Hervan, E.M., Mohammadi, S.A., Moradi, F., Tabatabaee, M.T., et al. (2015). QTL mapping of yield and yield components under normal and salt-stress conditions in bread wheat (*Triticum aestivum* L.). Plant Mol. Biol. Rep. 33(1), 102–120. doi: 10.1007/s11105-014-0726-0.

Bancal, P. (2008). Positive contribution of stem growth to grain number per spike in wheat. Field Crop Res. 105(1), 27–39. doi: 10.1016/j.fcr.2007.06.008.

Boden, S.A., Cavanagh, C., Cullis, B.R., Ramm, K., Greenwood, J., Jean Finnegan, E., et al. (2015). *Ppd-1* is a key regulator of inflorescence architecture and paired spikelet development in wheat. Nat. Plants 1(2), 14016. doi: 10.1038/nplants.2014.16.

Bridgemohan, P., and Bridgemohan, R.S.H. (2014). Evaluation of anti-lodging plant growth regulators on the growth and development of rice (*Oryza sativa*). J Cereals Oilseeds 5, 12–16. doi: 10.5897/JCO14.0128.

Cavanagh, C.R., Chao, S., Wang, S., Huang, B.E., Stephen, S., Kiani, S., et al. (2013). Genome-wide comparative diversity uncovers multiple targets of selection for improvement in hexaploid wheat landraces and cultivars. Proc. Natl. Acad. Sci. U. S. A. 110(20), 8057–8062. doi: 10.1073/pnas.1217133110.

Chae, K., Isaacs, C.G., Reeves, P.H., Maloney, G.S., Muday, G.K., Nagpal, P., et al. (2012). *Arabidopsis* SMALL AUXIN UP RNA63 promotes hypocotyl and stamen filament elongation. Plant J. 71(4), 684–697. doi: 10.1111/j.1365-313X.2012.05024.x.

Chen, Z., Cheng, X., Chai, L., Wang, Z., Du, D., Wang, Z., et al. (2020). Pleiotropic QTL influencing spikelet number and heading date in common wheat (*Triticum aestivum* L.). Theor. Appl. Genet. 133(6), 1825–1838. doi: 10.1007/s00122-020-03556-6.

Crossa, J., Jarquin, D., Franco, J., Perezrodriguez, P., Burgueno, J., Saintpierre, C., et al. (2016). Genomic prediction of gene bank wheat landraces. G3-Genes Genom. Genet. 6(7), 1819–1834. doi: 10.1534/g3.116.029637.

Cui, F., Ding, A., Li, J., Zhao, C., Wang, L., Wang, X., et al. (2012). QTL detection of seven spike-related traits and their genetic correlations in wheat using two related RIL populations. Euphytica 186(1), 177–192. doi: 10.1007/s10681-011-0550-7.

Deng, Z., Cui, Y., Han, Q., Fang, W., Li, J., and Tian, J. (2017). Discovery of consistent QTLs of wheat spike-related traits under nitrogen treatment at different development stages. Front. Plant Sci. 8, 2120. doi: 10.3389/fpls.2017.02120.

Dixon, L.E., Greenwood, J.R., Bencivenga, S., Zhang, P., Cockram, J., Mellers, G., et al. (2018). *TEOSINTE BRANCHED1* regulates inflorescence architecture and development in bread wheat (*Triticum aestivum*). Plant Cell 30(3), 563–581. doi: 10.1105/tpc.17.00961.

Dobrovolskaya, O., Pont, C., Sibout, R., Martinek, P., Badaeva, E., Murat, F., et al. (2015). *FRIZZY PANICLE* drives supernumerary spikelets in bread wheat. Plant Physiol. 167, 189–199. doi: 10.1104/pp.114.250043.

El Haddad, N., Kabbaj, H., Zaïm, M., El Hassouni, K., Tidiane Sall, A., Azouz, M., et al. (2021). Crop wild relatives in durum wheat breeding: Drift or thrift? Crop Sci. 61(1), 37–54. doi: 10.1002/csc2.20223.

Faris, J.D., Zhang, Q., Chao, S., Zhang, Z., and Xu, S.S. (2014a). Analysis of agronomic and domestication traits in a durum × cultivated emmer wheat population using a high-density single nucleotide polymorphism-based linkage map. Theor. Appl. Genet. 127, 2333–2348. doi: 10.1007/s00122-014-2380-1.

Faris, J.D., Zhang, Z., Garvin, D.F., and Xu, S.S. (2014b). Molecular and comparative mapping of genes governing spike compactness from wild emmer wheat. Mol. Genet. Genomics 289(4), 641–651. doi: 10.1007/s00438-014-0836-2.

Gao, X., Wang, N., Wang, X., and Zhang, X. (2019). Architecture of wheat inflorescence: insights from rice. Trends Plant Sci. 24, 802–809. doi: 10.1016/j.tplants.2019.06.002.

Hou, K., Wu, W., and Gan, S. (2013). *SAUR36*, a small auxin up RNA gene, is involved in the promotion of leaf senescence in *Arabidopsis*. Plant Physiol 161(2), 1002–1009. doi: 10.1104/pp.112.212787.

Kamran, A., Iqbal, M., and Spaner, D. (2014). Flowering time in wheat (*Triticum aestivum* L.): a key factor for global adaptability. Euphytica 197, 1–26. doi: 10.1007/s10681-014-1075-7.

Klymiuk, V., Yaniv, E., Huang, L., Raats, D., Fatiukha, A., Chen, S., et al. (2018). Cloning of the wheat *Yr15* resistance gene sheds light on the plant tandem kinase-pseudokinase family. Nat. Commun. 9(1), 3735. doi: 10.1038/s41467-018-06138-9.

Kumar, N., Kulwal, P.L., Balyan, H.S., and Gupta, P.K. (2007). QTL mapping for yield and yield contributing traits in two mapping populations of bread wheat. Mol. Breed. 19(2), 163–177. doi: 10.1007/s11032-006-9056-8.

Kuzay, S., Xu, Y., Zhang, J., Katz, A., Pearce, S., Su, Z., et al. (2019). Identification of a candidate gene for a QTL for spikelet number per spike on wheat chromosome arm 7AL by high-resolution genetic mapping. Theor. Appl. Genet. 132 (9), 2689–2705. doi: 10.1007/s00122-019-03382-5.

Li, C., Tang, H., Luo, W., Zhang, X., Mu, Y., Deng, M., et al. (2020). A novel, validated, and plant height-independent QTL for spike extension length is associated with yield-related traits in wheat. Theor. Appl. Genet. 133, 3381–3393. doi: 10.1007/s00122-020-03675-0.

Li, S., Jia, J., Wei, X., Zhang, X., Li, L., Chen, H., et al. (2007). A intervarietal genetic map and QTL analysis for yield traits in wheat. Mol. Breed. 20(2), 167–178. doi: 10.1007/s11032-007-9080-3.

Li, T., Deng, G., Tang, Y., Su, Y., Wang, J., Cheng, J., et al. (2021). Identification and validation of a novel locus controlling spikelet number in bread wheat (*Triticum aestivum* L.). Front. Plant Sci. 12, 611106. doi: 10.3389/fpls.2021.611106.

Lin, Y., Jiang, X., Hu, H., Zhou, K., Wang, Q., Yu, S., et al. (2021). QTL mapping for grain number per spikelet in wheat using a high-density genetic map. Crop J. doi: 10.1016/j.cj.2020.12.006.

Liu, D., Yen, C., Yang, J., Zheng, Y., and Lan, X. (1999). The chromosomal locations of high crossability genes in tetraploid wheat *Triticum turgidum* L. cv. Ailanmai native to Sichuan, China. Euphytica 108(2), 79–82. doi: 10.1023/A:1003691925501.

Liu, H., Ma, J., Tu, Y., Zhu, J., Ding, P., Liu, J., et al. (2020a). Several stably expressed QTL for spike density of common wheat (*Triticum aestivum*) in multiple environments. Plant Breed. 139, 284–294. doi: 10.1111/pbr.12782.

Liu, J., Luo, W., Qin, N., Ding, P., Zhang, H., Yang, C., et al. (2018). A 55 K SNP array-based genetic map and its utilization in QTL mapping for productive tiller number in common wheat. Theor. Appl. Genet. 131(11), 2439–2450. doi: 10.1007/s00122-018-3164-9.

Liu, J., Tang, H., Qu, X., Liu, H., Li, C., Tu, Y., et al. (2020b). A novel, major, and validated QTL for the effective tiller number located on chromosome arm 1BL in bread wheat. Plant Mol. Biol. 104(1), 173–185. doi: 10.1007/s11103-020-01035-6.

Lopes, M.S., Ibrahim, E.B., Baenziger, P.S., Sukhwinder, S., Conxita, R., Kursad, O., et al. (2015). Exploiting genetic diversity from landraces in wheat breeding for adaptation to climate change. J. Exp. Bot. 66(12), 3477–3486. doi: 10.1093/jxb/erv122.

Ma, J., Ding, P., Liu, J., Li, T., Zou, Y., Habib, A., et al. (2019a). Identification and validation of a major and stably expressed QTL for spikelet number per spike in bread wheat. Theor. Appl. Genet. 132(11), 3155–3167. doi: 10.1007/s00122-019-03415-z.

Ma, J., Qin, N., Cai, B., Chen, G., Ding, P., Zhang, H., et al. (2019b). Identification and validation of a novel major QTL for all-stage stripe rust resistance on 1BL in the winter wheat line 20828. Theor. Appl. Genet. 132(5), 1363–1373. doi: 10.1007/s00122-019-03283-7.

Ma, J., Tu, Y., Zhu, J., Luo, W., Liu, H., Li, C., et al. (2020). Flag leaf size and posture of bread wheat: genetic dissection, QTL validation and their relationships with yield-related traits. Theor. Appl. Genet. 133(1), 297–315. doi: 10.1007/s00122-019-03458-2.

Maccaferri, M., Harris, N.S., Twardziok, S.O., Pasam, R.K., Gundlach, H., Spannagl, M., et al. (2019). Durum wheat genome highlights past domestication signatures and future improvement targets. Nat. Genet. 51(5), 885–895. doi: 10.1038/s41588-019-0381-3.

Manickavelu, A., Kawaura, K., Imamura, H., Mori, M., and Ogihara, Y. (2011). Molecular mapping of quantitative trait loci for domestication traits and β-glucan content in a wheat recombinant inbred line population. Euphytica 177(2), 179–190. doi: 10.1007/s10681-010-0217-9.

Meng, L., Li, H., Zhang, L., and Wang, J. (2015). QTL IciMapping: Integrated software for genetic linkage map construction and quantitative trait locus mapping in biparental populations. Crop J. 3(3), 269–283. doi: 10.1016/j.cj.2015.01.001.

Mohler, V., Lukman, R., Ortiz-Islas, S., William, M., Worland, A.J., van Beem, J., et al. (2004). Genetic and physical mapping of photoperiod insensitive gene *Ppd-B1* in common wheat. Euphytica 138(1), 33–40. doi: 10.1023/B:EUPH.0000047056.58938.76.

Peng, J., Korol, A., Fahima, T., Roder, M., Li, Y., and Nevo, E. (2001). QTLs for agronomic traits in tetraploid wild emmer wheat, *Triticum dicoccoides*. J. Sichuan Agric. Univ. 19, 317–334. doi: 10.16036/j.issn.1000-2650.2001.04.005.

Peng, J., Ronin, Y.I., Fahima, T., Roder, M.S., Li, Y., Nevo, E., et al. (2003). Domestication quantitative trait loci in *Triticum dicoccoides*, the progenitor of wheat. P. Natl. Acad. Sci. USA 100(5), 2489–2494. doi: 10.1073/pnas.252763199.

Quintero, A., Molero, G., Reynolds, M.P., and Calderini, D.F. (2018). Trade-off between grain weight and grain number in wheat depends on GxE interaction: A case study of an elite CIMMYT panel (CIMCOG). Eur. J. Agron. 92, 17–29. doi: 10.1016/j.eja.2017.09.007.

Ren, T., Hu, Y., Tang, Y., Li, C., Yan, B., Ren, Z., et al. (2018). Utilization of a wheat55K SNP array for mapping of major QTL for temporal expression of the tiller number. Front. Plant Sci. 9, 333. doi: 10.3389/fpls.2018.00333.

Rothwell, W. (2019). Origin of Perl. In: Beginning Perl Programming. Apress, Berkeley, CA. doi: 10.1007/978-1-4842-5055-6_1.

Sakuma, S., Golan, G., Guo, Z., Ogawa, T., Tagiri, A., Sugimoto, K., et al. (2019). Unleashing floret fertility in wheat through the mutation of a homeobox gene. Proc. Natl. Acad. Sci. U. S. A. 116, 5182–5187. doi: 10.1073/pnas.1815465116.

Shang, Q., Zhang, D., Li, R., Wang, K., Cheng, Z., Zhou, Z., et al. (2020). Mapping quantitative trait loci associated with stem-related traits in maize (*Zea mays* L.). Plant Mol. Biol. 104, 583–595. doi: 10.1007/s11103-020-01062-3.

Sharma, J.S., Running, K.L.D., Xu, S.S., Zhang, Q., Peters Haugrud, A.R., Sharma, S., et al. (2019). Genetic analysis of threshability and other spike traits in the evolution of cultivated emmer to fully domesticated durum wheat. Mol. Genet. Genomics 294(3), 757–771. doi: 10.1007/s00438-019-01544-0.

Shaw, L.M., Turner, A.S., Laurence, H., Simon, G., Laurie, D.A., and Somers, D.E. (2013). Mutant alleles of *Photoperiod-1* in wheat (*Triticum aestivum* L.) that confer a late flowering phenotype in long days. PLoS One 8(11), e79459. doi: 10.1371/journal.pone.0079459.

Sthapit, S.R., Marlowe, K., Covarrubias, D.C., Ruff, T.M., Eagle, J.D., McGinty, E.M., et al. (2020). Genetic diversity in historical and modern wheat varieties of the U.S. Pacific Northwest. Crop Sci. 60, 3175–3190. doi: 10.1002/csc2.20299.

Sun, C., Dong, Z., Zhao, L., Ren, Y., Zhang, N., and Chen, F. (2020). The Wheat 660K SNP array demonstrates great potential for marker-assisted selection in polyploid wheat. Plant Biotechnol. J. 18(6), 1354–1360. doi: 10.1111/pbi.13361.

Tan, C.-T., Yu, H., Yang, Y., Xu, X., Chen, M., Rudd, J.C., et al. (2017). Development and validation of KASP markers for the greenbug resistance gene *Gb7* and the Hessian fly resistance gene *H32* in wheat. Theor. Appl. Genet. 130(9), 1867–1884. doi: 10.1007/s00122-017-2930-4.

Tu, Y., Liu, H., Liu, J., Tang, H., Mu, Y., Deng, M., et al. (2021). QTL mapping and validation of bread wheat flag leaf morphology across multiple environments in different genetic backgrounds. Theor. Appl. Genet. 134(1), 261–278. doi: 10.1007/s00122-020-03695-w.

Ur Rehman, S., Wang, J., Chang, X., Zhang, X., Mao, X., and Jing, R. (2019). A wheat protein kinase gene *TaSnRK2.9*-5A associated with yield contributing traits. Theor. Appl. Genet. 132(4), 907–919. doi: 10.1007/s00122-018-3247-7.

Voorrips, R.E. (2002). MapChart: software for the graphical presentation of linkage maps and QTLs. J. Hered. 93(1), 77–78. doi: 10.1093/jhered/93.1.77.

Wang, Y., Lin, L., Ye, T., Lu, Y., Chen, X., and Wu, Y. (2013). The inhibitory effect of ABA on floral transition is mediated by ABI5 in Arabidopsis. J. Exp. Bot. 64(2), 675–684. doi: 10.1093/jxb/ers361.

Wang, Y., Yu, H., Tian, C., Sajjad, M., Gao, C., Tong, Y., et al. (2017). Transcriptome association identifies regulators of wheat spike architecture. Plant Physiol. 175(2), 746–757. doi: 10.1104/pp.17.00694.

Wu, X., Chang, X., and Jing, R. (2012). Genetic insight into yield-associated traits of wheat grown in multiple rain-fed environments. PloS One 7, e31249. doi: 10.1371/journal.pone.0031249.

Xie, W., and Nevo, E. (2008). Wild emmer: genetic resources, gene mapping and potential for wheat improvement. Euphytica 164(3), 603–614. doi: 10.1007/s10681-008-9703-8.

Yao, H., Xie, Q., Xue, S., Luo, J., Lu, J., Kong, Z., et al. (2019). *HL2* on chromosome 7D of wheat (*Triticum aestivum* L.) regulates both head length and spikelet number. Theor. Appl. Genet. 132(6), 1789–1797. doi: 10.1007/s00122-019-03315-2.

Yao, J., Ren, L., Zhang, P., Yang, X., and Zhou, M. (2011). Genetic and correlation analysis of plant height and its components in wheat. J. Triticeae Crop 31, 604–610 (in Chinese with English abstract).

Yu, J., Zhao, Y., Ding, M., Yu, Z., Jiang, Y., Ma, W., et al. (2020). Wild emmer chromosome arm substitution lines: Useful resources for wheat genetic study and breeding. Crop Sci. 60, 1761–1769. doi: 10.1002/csc2.20022.

Zaïm, M., El Hassouni, K., Gamba, F., Filali-Maltouf, A., Belkadi, B., Sourour, A., et al. (2017). Wide crosses of durum wheat (*Triticum durum* Desf.) reveal good disease resistance, yield stability, and industrial quality across Mediterranean sites. Field Crop Res. 214, 219–227. doi: 10.1016/j.fcr.2017.09.007.

Zhu, T., Wang, L., Rimbert, H., Rodriguez, J.C., Deal, K.R., De Oliveira, R., et al. (2021). Optical maps refine the bread wheat *Triticum aestivum* cv. Chinese Spring genome assembly. Plant J. doi: 10.1111/tpj.15289.

Zhu, T., Wang, L., Rodriguez, J.C., Deal, K.R., Avni, R., Distelfeld, A., et al. (2019). Improved Genome Sequence of Wild Emmer Wheat Zavitan with the Aid of Optical Maps. G3: Genes. Genom. Genet. 9(3), 619–624. doi: 10.1534/g3.118.200902.

