## Supplementary figures and images for "The 55K SNP-Based Exploration of QTL for Spikelet Number Per Spike in a Tetraploid Wheat (*Triticum turgidum* L.) Population: Chinese Landrace ‘Ailanmai’ × Wild Emmer"

### Supplementary Figure 1

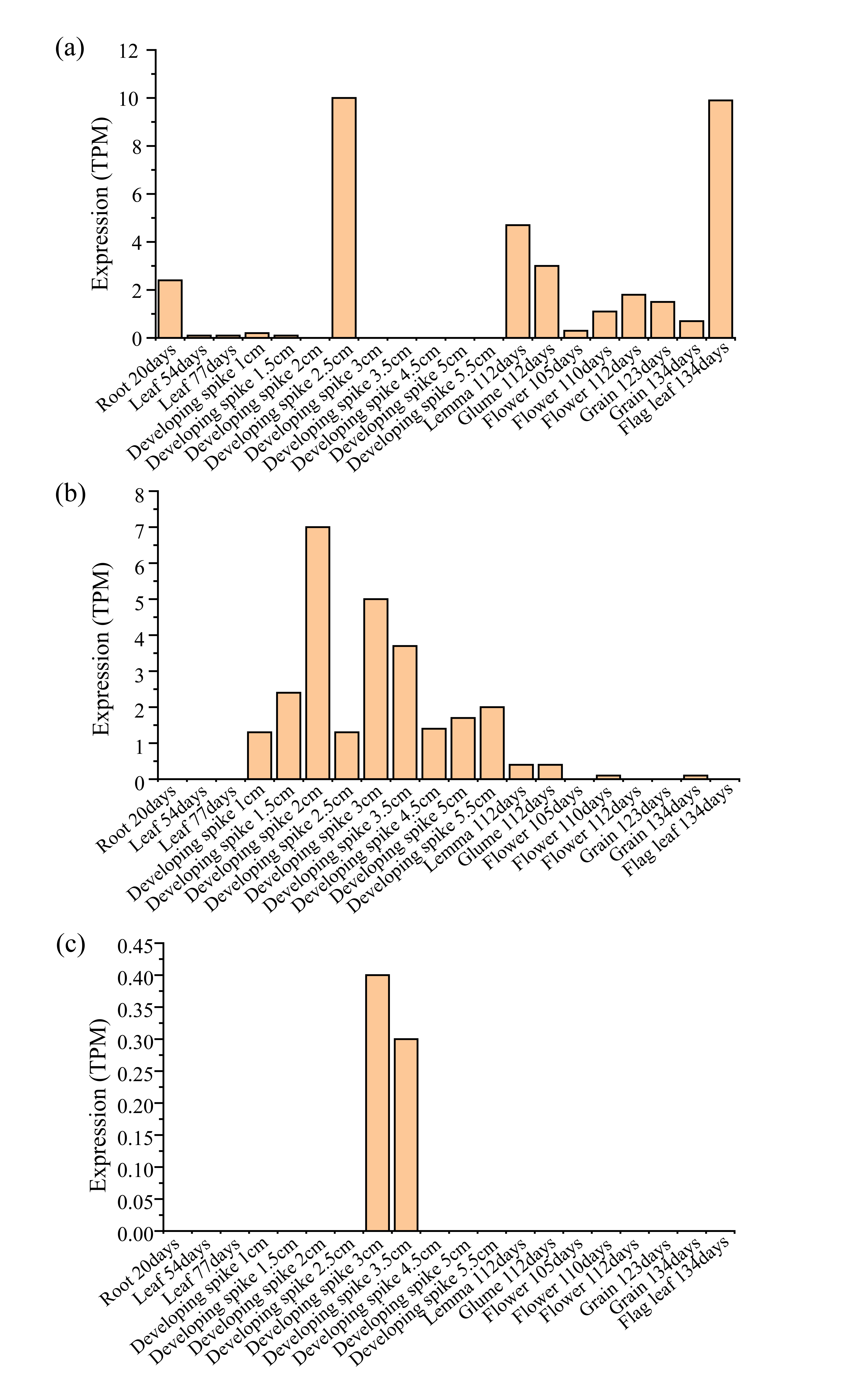
